# Mesophilic enzyme function at high temperature: molecular dynamics of hyperthermophilic and mesophilic pyrophosphatases

**DOI:** 10.1101/2020.03.05.979179

**Authors:** Rupesh Agarwal, Utsab R. Shrestha, Xiang-Qiang Chu, Loukas Petridis, Jeremy C. Smith

## Abstract

The mesophilic inorganic pyrophosphatase from *Escherichia coli* (*EcPPase*) retains function at 353 K, the physiological temperature of hyperthermophilic *Thermoccoccus thioreducens*, whereas, the homolog protein from the hyperthermophilic organism (*Tt*PPase) cannot function at room temperature. To explain this asymmetric behavior, we examined structural and dynamical properties of the two proteins using molecular dynamics simulations. The global flexibility of *Tt*PPase is significantly higher than its mesophilic homolog at all tested temperature/pressure conditions. However, at 353 K, *Ec*PPase reduces its solvent-exposed surface area and increases subunit compaction while maintaining flexibility in its catalytic pocket. In contrast, *Tt*PPase lacks this adaptability and has increased rigidity and reduced protein:water interactions in its catalytic pocket at room temperature, providing a plausible explanation for its inactivity near room temperature.

## Introduction

The enzymatic activity of proteins from hyperthermophilic microorganisms thriving in extreme conditions has been an active area of research for several decades^1,2^. These microbes have an optimal temperature range for their growth and survival of about 80°C - 100°C ^3^. In addition to extreme temperatures, these microorganisms can also withstand high hydrostatic pressures ranging from 60 to 100 MPa (the atmospheric pressure at sea-level is 0.1 MPa)^1,4^. Since most mesophilic proteins denature under such high temperatures^5^ and pressures^6^, it is intriguing to examine how proteins from hyperthermophilic organisms retain activity. Previous studies have suggested structural characteristics^7–9^ of proteins that enable these extremophilic microbes to thrive in severe conditions. However, there appears to be no universal adaptive mechanism, but rather a complex combination of different factors, which frequently differs according to the protein or protein family and is thus difficult to generalize^8,10^. Furthermore, the role of dynamic characteristics such as conformational stability^11 12^ and flexibility^13^, for protein adaptability is still not well understood^1,14^.

Recent studies have reported that the conformational sub-states of a protein are significantly perturbed by changes in temperature and pressure^13,15–19^. Temperature enhances the internal fluctuations of a protein^20^, and an optimum temperature may provide an appropriate balance of flexibility and rigidity required for function^13,21,22^. Temperatures higher than the optimum can lead to loss of function through unfolding or denaturation^23^. However in the case of hyperthermophilic proteins, high native-state flexibility can reduce their entropy of unfolding, thus increasing their melting temperature^24^. Similarly, high pressure conditions can cause a protein to become inactive by the collapse of its intra-protein cavities, giving rise to an unfolded state^25^. Pressure drives the reduction in the volume of a protein, which results in a negative entropy change of the system, which may destabilize the native state^26^. Nonetheless, there are exceptions, and the stability and/or activity of some proteins, such as a thermolysin^27^ and a hydrogenase from *Methanococcus jannaschii*^28^, have been shown to increase with pressure.

Overall, a dualistic picture of protein flexibility^24,29^ and rigidity^30,31^ has been recognized as a possible factor behind thermostability of thermophilic and hyperthermophilic proteins^13,19^. On the one hand, flexibility is required by a protein to function. On the other hand, flexible residues trigger protein unfolding due to their large thermal fluctuations at high temperature. Hence, rigidifying flexible residues may be an effective way to improve thermostability^30,31^. In addition to flexibility, oligomerization has been reported to be critically important for the stability of some proteins^32,33^ but the structure-based reasoning behind this stability is not understood^34^.

We compare here a particularly interesting pair of proteins - the inorganic pyrophosphatase (PPase) from *Thermococcus thioreducens* (*Tt*), a hyperthermophilic archaea found near hydrothermal vents of the Mid-Atlantic Ridge^35^, with a homolog from the mesophilic bacterium, *Escherichia coli* (*Ec*)^36^. PPase (EC 3.6.1.1) is a homo-hexameric enzyme (~120 kDa), which catalyzes the conversion of pyrophosphate into two phosphate ions. This conversion is important for many critical biochemical processes, such as the production of proteins, nucleic acid polymerization, and lipid metabolism^35^. Although *Tt*PPase and *Ec*PPase have ~60% sequence similarity and ~40% identity, and share similar oligomeric crystal structures (**Fig. S2**), the temperatures and pressures for their optimal enzymatic activities are very dissimilar. The catalytic activity of *Tt*PPase has been reported to be maximal at ~353 K but negligible (1.3% of the maximum activity in 10 mins) near room temperature and standard atmospheric pressure (1 bar or 0.1 MPa) ^37^. In contrast, the optimal conditions for *Ec*PPase have been shown to be room temperature (298 K) and standard atmospheric pressure. Interestingly, unlike other mesophilic proteins, *Ec*PPase retains up to 95 to 100% of its enzymatic activity at 353 K for about 10 minutes, after which the activity slowly decays^38^.

Most crystal structures of thermophilic proteins have been resolved at room temperature instead of at their native temperature or pressure conditions. Therefore, to obtain a comprehensive picture of protein structure, dynamics, and function, static structure determination (using X-ray and neutron crystallography) must be complemented with dynamic information using various spectroscopic techniques (e.g., neutron scattering^13,21^ or NMR^39,40^) and computer simulations^41,42^ under varying external conditions. Molecular dynamics (MD) simulation, which provides spatial and temporal information at atomic resolution^43^ has thus become one of the most powerful methods to explore the protein energy landscapes and their flexibility-function relationships^18,19,44^ at near-native conditions and is the technique applied here.

Here, we perform MD simulations to understand the effects of pressure and temperature on the structural and dynamic behavior of the two PPase homologs in their native and non-native environments. We find that *Tt*PPase possesses higher global flexibility at both native and non-native conditions than its mesophilic homolog (*Ec*PPase). However, this effect is not reflected locally in the catalytic pocket. Additionally, we determined that factors accompanying enzymatic activity of PPase are the number of hydrogen bonds and water molecules in its catalytic pocket. Furthermore, we provide potential factors behind the observed enzyme activity and/or adaptability of *Ec*PPase at high temperature and alternatively the inability of *Tt*PPase to do so at low temperature.

## Results

### I. Structural flexibility

The structural flexibility of proteins is associated with many biological functions such as catalytic activity, substrate binding, and molecular recognition^45,46^. To understand the effect of temperature and pressure on the protein global flexibility, we calculated the average mean-square displacement (MSD) of the Cα atoms. The MSD is often used to compare the stability of thermophilic and mesophilic protein homologs^18,19^. Overall, the MSD results show that *Tt*PPase has higher flexibility compared to *Ec*PPase at all the conditions studied here **(Fig. 1)**. This is therefore an example of a thermophilic protein for which increased rigidity is not associated with thermostability.

**Figure 1:**
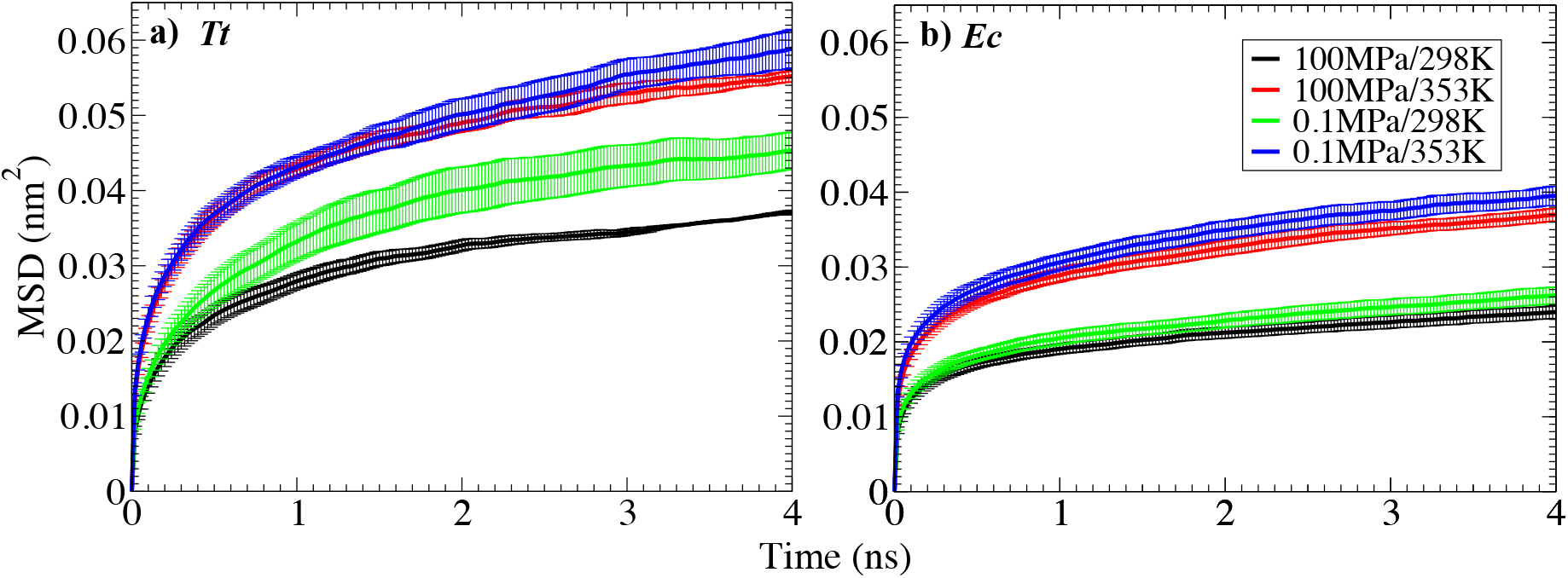
Mean square displacement (MSD) of C-alpha atoms for (a) hexameric *Tt*PPase and (b) hexameric *Ec*PPase. Higher values of MSD for *Tt*PPase at all conditions indicate higher structural flexibility than *Ec*PPase. Error bars shown in this and the subsequent figures are the standard error of the mean, and those not immediately visible are at most the size of the symbol.

For both the homologs, a two-fold increase in MSD is observed at 353 K relative to 298 K. Unlike temperature, the effect of pressure was less pronounced, an increase in pressure causing a small reduction in MSD for *Tt*PPase, and no change for *Ec*PPase. Interestingly, the MSD values for both the monomers showed an effect of pressure at 353 K, contrary to the respective hexameric forms (**Fig. 1 and S5**). These results suggest that oligomerization may confer stability to increased pressure at high temperature.

Flexibility is often correlated with activity or stability of mesophilic and thermophilic homologs^45,46^. However, although at 298 K *Tt*PPase is more flexible than *Ec*PPase, the activity of *Tt*PPase is negligible near room temperature^37^. Further, even though the flexibility of *Ec*PPase at 298 K is smaller than that at 353 K, *Ec*PPase is catalytically active at both temperatures. Hence, the global flexibility of the PPase does not follow the observed functional properties.

### II. Compactness

Compactness is another common adaptive mechanism of thermophilic proteins^34,47^. To characterize the compactness of the hexameric structures over the course of the simulations, we calculated the solvent accessible surface area (SASA) of the proteins, which is inversely proportional to the number of native contacts^48^. The density plots of the SASA show that *Ec*PPase has larger solvent accessibility than *Tt*PPase, irrespective of the temperature and pressure conditions studied here (**Fig. 2a and S1b**). A similar difference is found in the hexameric crystal structures (*Tt*PPase: 374 nm^2^ and *Ec*PPase: 393 nm^2^; **Table S3**). However, the solvent accessibility of *Ec*PPase is reduced at 353 K compared to 298 K, whereas that of *Tt*PPase remains unaffected by the change in temperature. Pressure did not significantly affect the SASA of either homolog. The SASA probability distribution for monomeric structures showed a similar change in compactness as observed in the hexameric form (**Fig S4a**).

**Figure 2:**
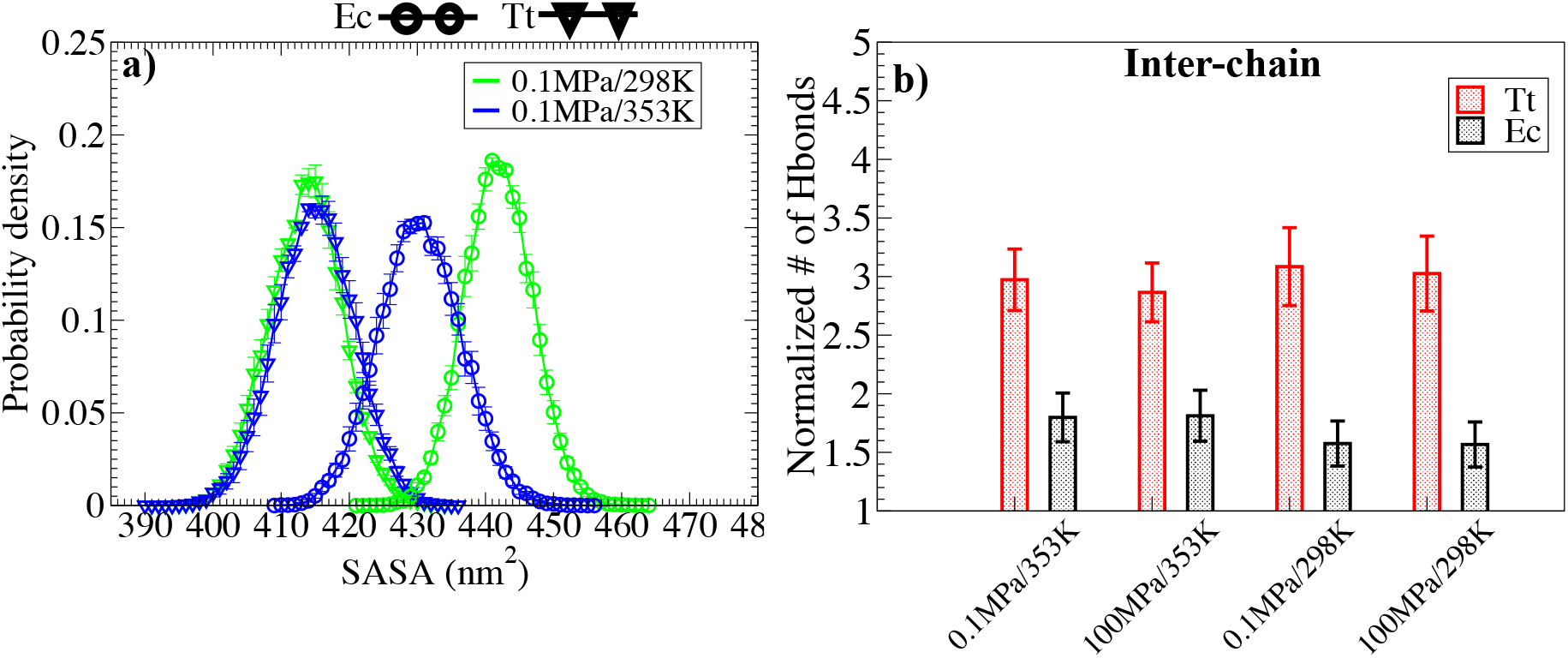
a) Probability distribution for solvent accessible surface area (SASA) of the hexameric forms for *Tt*PPase (triangle) and *Ec*PPase (circle); b) The normalized number of interchain hydrogen bonds.

To further quantify compactness, we calculated the number of intra- and inter-chain hydrogen bonds (HBs) of the hexameric form. The normalized mean number of inter-chain HBs was nearly double for *Tt*PPase than for *Ec*PPase and remained constant at all conditions (**Fig. 2b**), confirming the closer packing of the subunits in *Tt*PPase than in *Ec*PPase. The mean number of intra-chain HBs did not change greatly between the two homologs (**Fig. S1a**) at all pressure and temperature conditions. These results suggest that the higher compactness of the *Tt*PPase hexamer is due to compactness between the subunits and also within each monomer.

### III. Protein cavities

Intra-protein cavities have been recognized to be important for the stability and function of proteins^49,50^, and water molecules buried within these cavities have been reported to be influential in temperature and pressure-mediated unfolding^25,51^. We calculated the number of water molecules enclosed in the completely buried cavities of both proteins. This was found to be significantly higher for *Ec*PPase than *Tt*PPase (**Fig. 3**) and not significantly affected by the temperature and pressure. These results are consistent with the inter-chain Hbond results discussed earlier and with calculations that show that the crystal structure of *Tt*PPase has smaller buried cavity volume (303 water molecules) than *Ec*PPase (427 water molecules) (**Table S3**).

**Figure 3:**
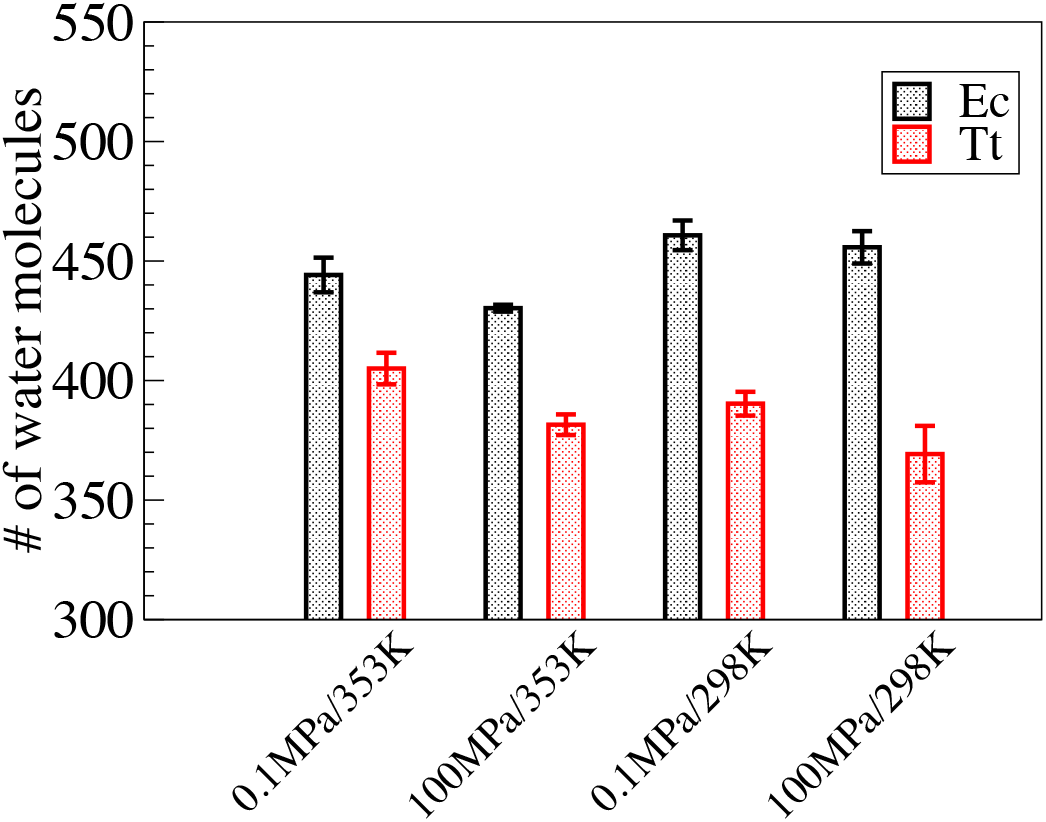
Number of water molecules in completely buried cavities of hexameric form at different temperature and pressure conditions. *Tt*PPase (black shaded bars) vs. *Ec*PPase (red shaded bars).

### IV. Flexibility in the catalytic pocket and its interaction with water

The precise 3D arrangement of residues and their interaction with water molecules in an enzyme’s catalytic pocket determines the function ^52,53^. Previous studies have identified conserved residues in the catalytic pocket of PPase enzymes (**Table S1 and Figs. S2a and S2b**) that are critical for catalysis and are also required for coordinating the divalent ions^54,55^. The catalytically inactive *Tt*PPase at 298 K has ~60% more HBs (~15 HBs i.e., 2.5 per monomer) between the catalytic residues than the catalytically-active forms of PPasse (*Tt*PPase at 353 K and *Ec*PPase at 298 and 353 K) (**Figs. 4 a, c**). These results, for the hexameric form were also found in the simulations of the monomeric form (**Fig. S4b)**. We did not observe any effect of pressure on the number of HBs between the residues in the catalytic pocket **(Figs. 4 a, c)**.

**Figure 4:**
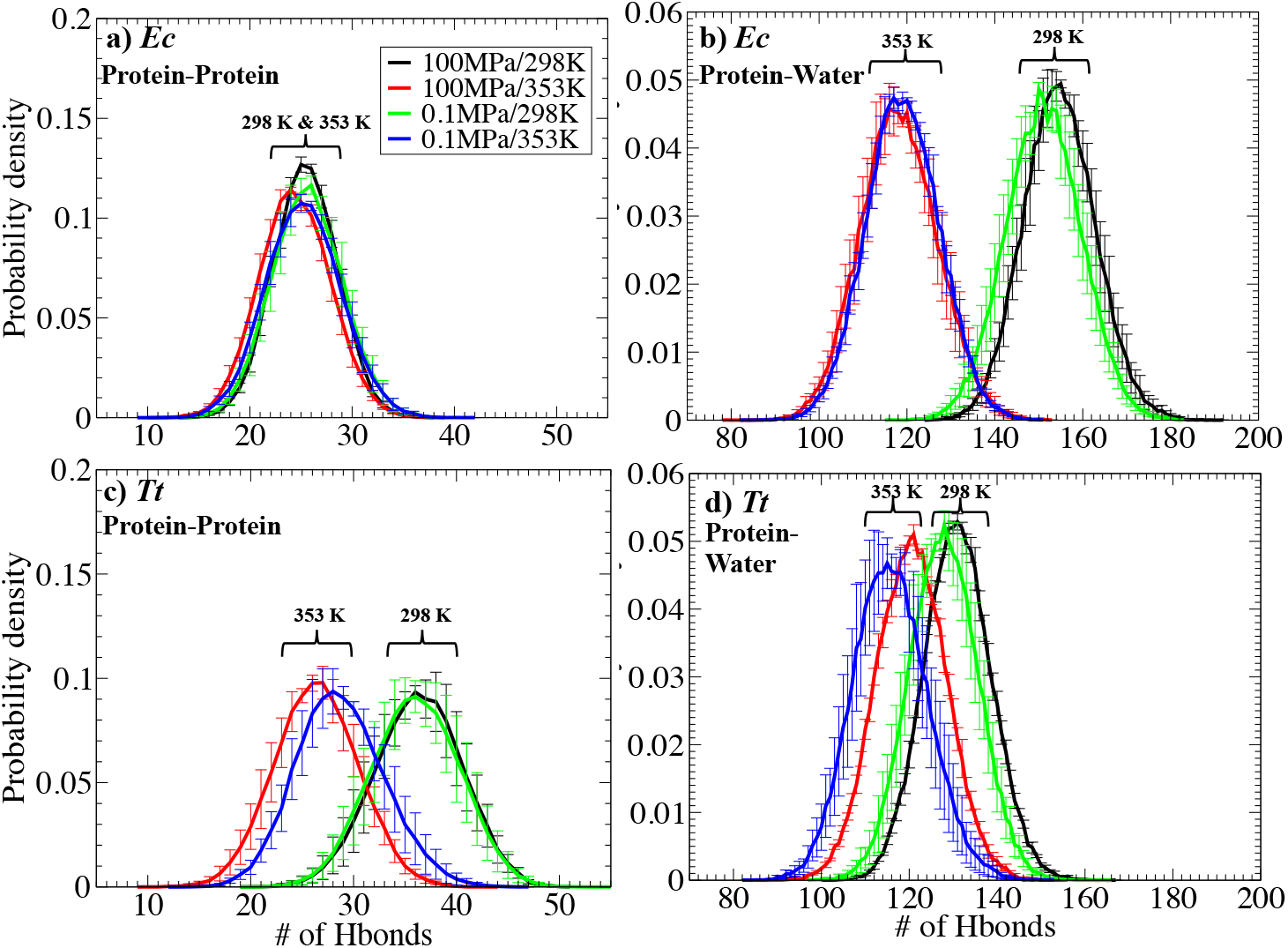
Probability distributions of the number of hydrogen bonds between the residues in the catalytic pocket: hexameric *Ec*PPase (a) and hexameric *Tt*PPase (c). Probability distributions of the number of hydrogen bonds between the catalytic pocket residues and water molecules in the catalytic pocket: hexameric *Ec*PPase (b) and hexameric *Tt*PPase (b).

Water molecules in the catalytic pockets can be critical for enzymatic function^56,57,58^. Interestingly, the catalytic pockets of both homologs have ~110 protein:water HBs at 353 K, consistent with the observed catalytic activity of *Ec*PPase at this elevated temperature^38^. However, at 298 K, *Ec*PPase significantly increases the number of protein:water HBs, to around ~160 HBs, whereas *Tt*PPase does this to a much lesser degree, to ~130 HBs **(Figs. 4 b, d)**.

We also quantified the flexibility and solvent exposure of the catalytic pocket by calculating the root mean square fluctuation (RMSF) of these conserved residues. The RMSF of these residues in *Ec*PPase varies weakly with temperature, whereas *Tt*PPase has significantly lower fluctuations at 298 K compared to 353 K **(Fig. S3)**. *Ec*PPase therefore maintains its local flexibility of the catalytic pocket at both temperatures whereas *Tt*PPase becomes rigid at the lower temperature.

## Discussion and Conclusions

In this work, we compare structural and dynamic properties of hyperthermophilic and mesophilic PPases using MD simulations mimicking deep-sea and ambient conditions. The results indicate that *Tt*PPase has been designed to function at high temperatures with a more compact structure and reduced number of intra-protein hydrogen bonds in the catalytic pocket. However, *Tt*PPase does not maintain both of these properties at room temperature, where it cannot catalyze the enzymatic reaction. Interestingly, we found *Ec*PPase adapts and retains its activity at high temperature by incorporating similar strategies used by *Tt*PPase to function at high temperature: maintaining hydrogen bonds in the catalytic pocket and increasing its compactness.

The overall structural flexibility of a protein is sometimes assumed to be relevant for its enzymatic activity^45^ and the MSD is often used to quantify this. A general framework for understanding the stability and function of hyperthermophilic proteins in their native conditions has been proposed based upon the hypothesis that enhanced rigidity underlies increased thermal stability^24^. However, other experimental and computational studies have reported that hyperthermophilic proteins have larger conformational flexibility than their mesophilic homologs^19,59^. Here, we also observe higher overall flexibility of *Tt*PPase than *Ec*PPase at all temperature/pressure conditions. Indeed, even at room temperature, where *Tt*PPase is enzymatically inactive, it is more flexible than *Ec*PPase. Thus, differences in the overall flexibility of PPase are not directly associated with differences its enzymatic activity.

Although the crystal structures of both homologs were not resolved at their native conditions, the RMSD between them is small (0.18 nm) and both homologs have similar radii of gyration (2.9 nm). However, the buried cavity size and solvent accessibility of the crystal structure of *Ec*PPase are greater than for *Tt*PPase. Likewise, from the simulations the solvent-accessible surface area and number of inter-chain Hbonds (both of which are related to compactness) of *Tt*PPase indicate that it is more compact than *Ec*PPase at all the conditions investigated here. The main contributing factors to the compactness of *Tt*PPase are inter-subunit compaction as shown by increased inter-chain H-bonding, together with the reduced solvent-exposed surface area of each monomer as shown by the simulations of the monomeric forms. These results are not surprising since compactness is a known adaptive factor in common between many thermophilic proteins.

Moreover, a decrease in the number of water molecules in the buried cavities of *Tt*PPase compared to *Ec*PPase was observed at all conditions. These results are consistent with previous work^60^ which showed that proteins that are active at extremely low temperatures (psychrophilic proteins) have a comparatively larger average cavity size. Similarly, a reduction in buried cavity volume has been reported as an adaptation (thermostability) mechanism at high-temperature conditions because large water-filled cavities are known to be a driving factor for protein denaturation at high temperature and pressure^61^.

Interestingly, *Ec*PPase exhibited a change in its compactness with a change in temperature, becoming more compact at higher temperature than at its natural temperature. In contrast, the compactness of *Tt*PPase remains the same at room and high temperatures. Additionally, we also studied the structure and dynamics of the catalytic pocket. Here, at 298 K we observed an increase in rigidity at 298 K compared to 353 K, as shown by the lower RMSF of catalytic pocket residues and a greatly increased number of H-Bonds between the catalytic pocket residues. In contrast, we observed similar catalytic pocket residue H-bonds in *Ec*PPase as for *Tt*PPase at 353 K, and this number remained unaltered upon change in temperature. This local flexibility (based upon the number of HBs and RMSF) of the catalytic pocket agrees with the existing “corresponding states” hypothesis, according to which a thermophilic protein has a more rigid catalytic pocket than its mesophilic homolog at room temperature^62^. However, both of these homologs should exhibit similar flexibility at their respective functional temperatures^62^. Furthermore, we see that the trend observed in global flexibility measured by the MSD is not reflected locally in the catalytic pocket.

The above picture is further supported by calculations of the number of H-bonds of the catalytic pocket residues with water molecules. In the case of *Ec*PPase, we observe a dramatic decrease in the number of Hbonds with water at 353 K when compared to 298 K, whereas there is a negligible change for *Tt*PPase. This result is consistent with the compactness results showing that *Ec*PPase has the capability to adapt to a high temperature whereas *Tt*PPase lacks the ability to adapt to low temperature environment.

Based on these results, we suggest that at lower temperatures, both of these homologs would need comparatively more water molecules in their catalytic pockets and a larger exposed surface area to function and that the opposite is needed at higher temperatures. *Ec*PPase is able to make the aforementioned changes to its structure and hence is able to adapt to high temperature/pressure but *Tt*PPase fails to do so at low temperature. Although the overall structure of *Tt*PPase barely responds to change in temperature/pressure, the catalytic pocket becomes more rigid at 293 K than at 353 K due to an increase in the number of intra-protein HBs. This rigidity may explain its enzymatic inactivity at room temperature. In contrast, *Ec*PPase adapts to high temperature by reducing solvent-exposed surface and adopting a more compact oligomeric structure. In addition, *Ec*PPase preserves the number of hydrogen bonds within the residues in the catalytic pocket at all the conditions and reduces its interactions with water molecules at higher temperature. This provides a possible explanation behind its activity even at a high temperature.

We observed an effect of pressure on the MSD of monomeric form, but not for hexameric form. However, the SASA and the catalytic pocket behavior of the monomeric and hexameric forms are relatively unaffected. Consequently, in the scope of our study, we suggest that the selective pressure behind oligomerization in the case of PPase remains unclear.

Finally, our results provide a structural and dynamic basis for the experimentally observed activity of *Ec*PPase at both high and room temperature. Such an intriguing ability of adaptation of mesophilic PPase could be possibly explained by existing evolutionary theory since hyperthermophilic archaea are thought to be a universal ancestor^63^. Indeed, recent work^64^ has suggested that thermophilic proteins (isolated from thermophilic archaea *Pyrococcus furiosus*) were ‘‘*de novo*’’ designed in a hot environment and then used a “structure-based mechanism” to adapt to a mesophilic environment on recolonization. In the structure-based mechanism adaptation to high temperatures is through compaction, which here is mostly inter-subunit, with the monomer structures relatively unchanged. Phylogenetic analyses^65^ show that *T. thioreducens (*also hyperthermophilic archaea) shares the same clade as *P. furiosus*; hence, PPase homologs studied here may have evolved according to the structure-based adaptation theory. Thus, *Ec*PPase may have been evolved from its ancestral counterpart *Tt*PPase to recolonize at room temperature and may therefore be able to adapt when introduced to a higher temperature. Moreover, since thermophiles do not live at room temperature, there is no selective need for high activity at room temperature.

## Methods

All-atom MD simulations of monomeric and hexameric *E. coli* PPase (*Ec*PPase) and *Thermococcus thioreducens* PPase (*Tt*PPase) were carried using GROMACS (version 2016.3) suite^66,67^. Analysis of the MD trajectories was performed using python package pytraj^68^ and GROMACS analysis tools. Hexameric *Ec*PPase (PDB ID 1I6T^36^) and *Tt*PPase (PDB ID 3Q5V^35^) were solvated in a TIP3P cubic box of water (~150,000 atoms). The crystal structures for both the proteins were resolved at the same conditions (298 K and 0.1 MPa) and are also in the same space group (H32). The monomeric structures for both the proteins were modeled by taking only one chain from the hexameric structures. These monomeric models were then solvated in a TIP3P cubic box (~36,000 atoms).

### Simulation details

All simulations were performed using the AMBER99SB potential^66^. Each protein was simulated at four different conditions: two temperatures: 298 K (ambient for *Ec*PPase) and 353 K (ambient for *Tt*PPase); and two pressures: 0.1 MPa (ambient for *Ec*PPase) and 100 MPa (ambient for *Tt*PPase). All systems were prepared in a three-step process: initial energy minimization, NVT equilibration and a NPT production run. Energy minimization was performed with the steepest-descent algorithm^66,67^ to a tolerance of 1000 kJ/ (mol·nm). NVT equilibration was performed for 1 ns at each temperature fixed with the V-Rescale thermostat^69^. Following this step, the NPT ensemble protocol was followed to generate production runs (500 nanoseconds for the hexamer and 1 microsecond for the monomers) with the V-Rescale thermostat^69^ used to maintain the required temperature and Parrinello-Rahman^70^ to maintain the pressure at 0.1 MPa or 100 MPa. Since the compressibility of water changes drastically with pressure, the compressibility of water was taken to be 4.5 × 10^−5^ bar ^−1^ at 0.1 MPa and 3.5 × 10^−5^ bar ^−1^ at 100 MPa, based on previous experimental work^71^. Particle Mesh Ewald^72^ was used for long-range electrostatics with a short-range electrostatics and van der Waals cutoff of 1 nm. Three independent runs were performed for each simulation (**Table S2**). The simulations were considered converged when the fluctuations in the root mean square deviation of C-alpha atoms (RMSD_Cα_) reached a plateau with time.

### Analysis details

Each of the metric was calculated from the last 300 ns of each simulation trajectory. The average value of each metric from 3 independent trajectories starting with different velocities has been reported, where the standard error of the mean has been used as the error bar.

The crystal structure RMSD comparison was performed using the SuperPose websever^73^. The number of hydrogen bonds was computed using the “hbond” utility in GROMACS using a donor−acceptor cutoff distance of 0.32 nm and a cutoff angle of 20°. The solvent accessible surface area (SASA) and volume were calculated using a 0.14 nm probe size for the whole protein. The volume of buried cavities was determined using the “trj_cavity” module^74^ in GROMACS. The volume of the completely buried cavities (CBC) was calculated with a 1.4 Å grid spacing (-spacing) and degree of buriedness of 6 Å. The number of water molecules in the cavities was estimated by dividing the total cavity volume by the volume of a buried water molecule near the protein surface (2.29 nm^3^) as reported in previous work^75^. Intra and interchain hydrogen bonds were calculated using the pytraj python package^68^. Inter chain hydrogen bonds were normalized by dividing it by the number of chains (n=6).

## Acknowledgments

We would like to thank Dr. Micholas Dean Smith and Dr. Omar Demerdash at ORNL for their valuable discussions. This research made use of the Compute and Data Environment for Science (CADES) computing resource at the Oak Ridge National Laboratory (ORNL), which is supported by the Office of Science of the U.S. Department of Energy under Contract No. DE-AC05-00OR22725. This research was supported by the Genomic Science Program, Office of Biological and Environmental Research, U. S. Department of Energy (DOE), under Contract FWP ERKP752 and of the National Energy Research Scientific Computing Center, a DOE Office of Science User Facility supported by the Office of Science of the U. S. Department of Energy under Contract No. DE-AC02-05CH11231.

