## Supplementary information for "Mesophilic enzyme function at high temperature: molecular dynamics of hyperthermophilic and mesophilic pyrophosphatases"

[1]UT/ORNL Center for Molecular Biophysics, Biosciences Division, Oak Ridge National Laboratory, Tennessee 37831-6309; [2]Graduate School of Genome Science and Technology, University of Tennessee, Knoxville, Tennessee 37996; [3]Department of Biochemistry and Cellular and Molecular Biology, University of Tennessee, Knoxville, Tennessee 37996; [4]Department of Nuclear Science and Technology, Graduate School of China Academy of Engineering Physics, Beijing 100193, China.

* co-corresponding authors

### SUPPLEMENTARY INFORMATION

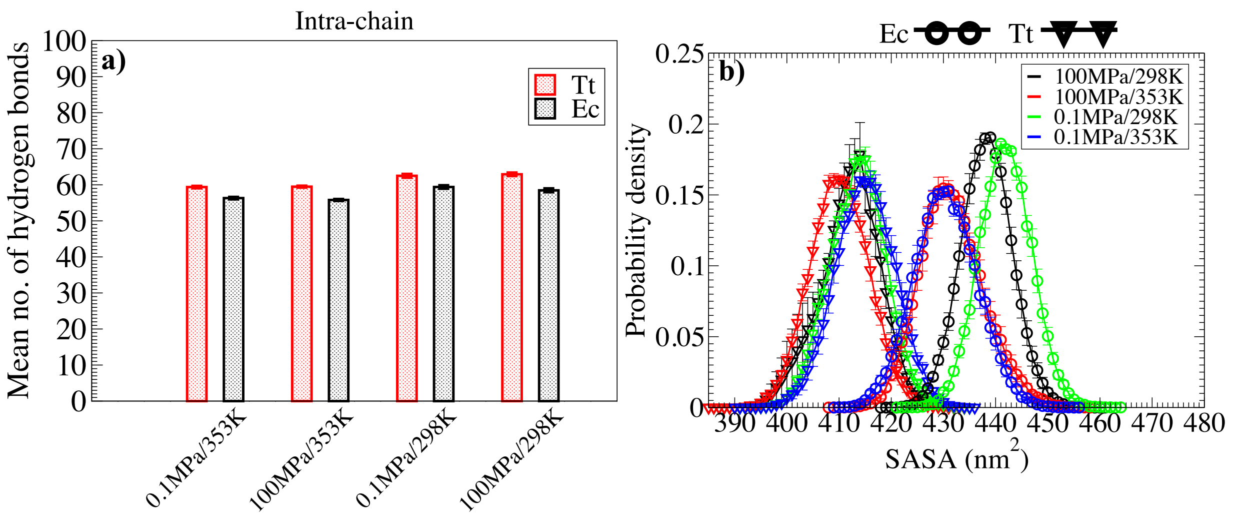


**Fig S1: a) Mean number of hydrogen bonds within each chain (intrachain) for hexameric *Tt*PPase and hexameric *Ec*PPase at different conditions; b) Probability distribution for solvent accessible surface area (SASA) of the hexameric forms for *Tt*PPase (triangle) and *Ec*PPase (circle); Error bars show the standard error of the mean (SEM), and those not immediately visible are the same size of the symbol.**

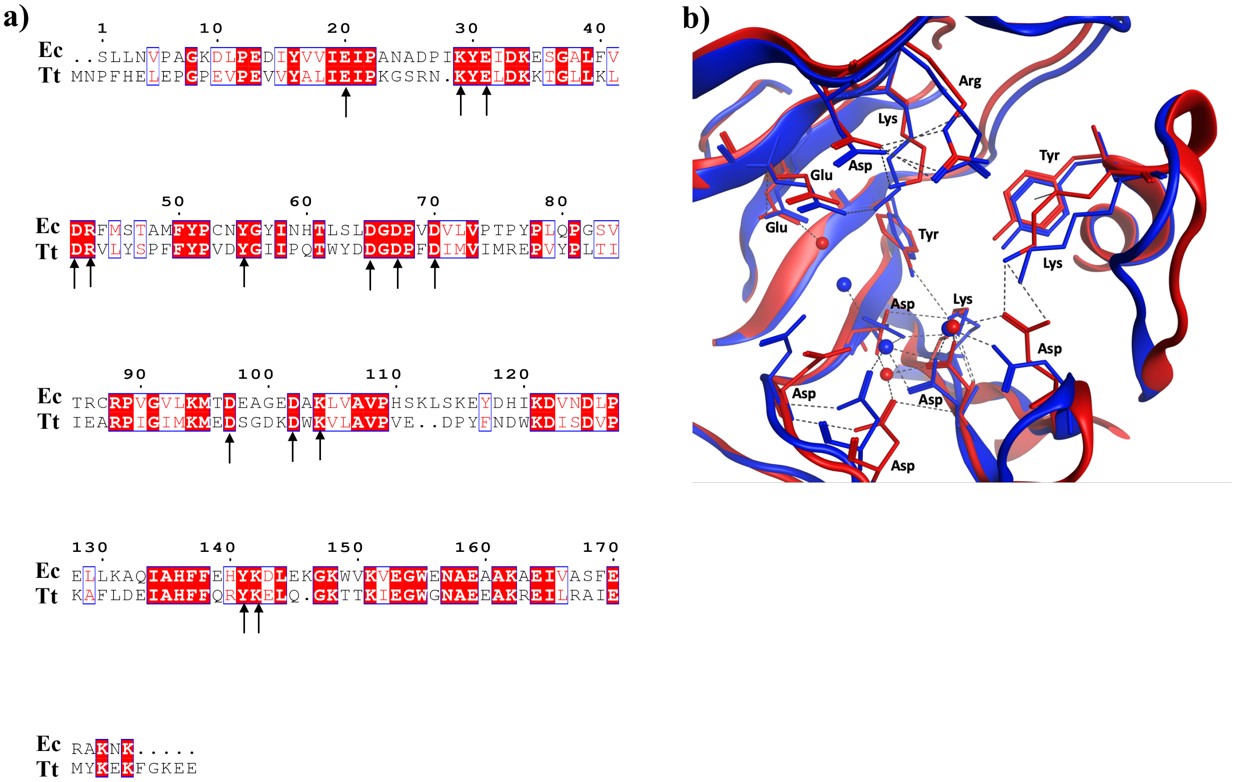


**Fig S2: a) Sequence alignment of *Ec*PPase (Ec) and *Tt*PPase (Tt) showing identical residues (in red box), similar residues (in white box) and conserved catalytic pocket residues (black arrow); b) Catalytic pocket of *Ec*PPase (red) and *Tt*PPase (blue) showing conserved critical residues (as sticks) and hydrogen bonds (black dotted line).**

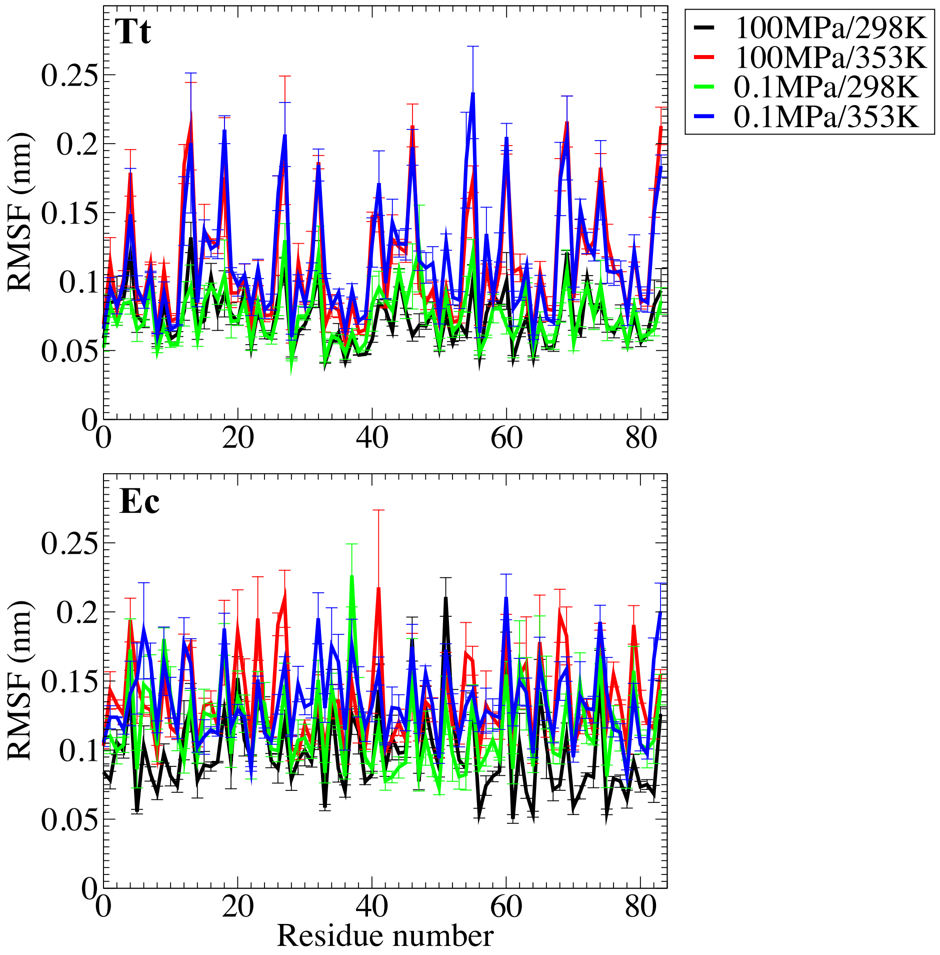


**Fig S3: Root mean square fluctuation (RMSF) of all atoms of catalytic pocket residues (averaged over per residue) for hexameric *Tt*PPase (top) and hexameric *Ec*PPase (bottom) at different conditions.**

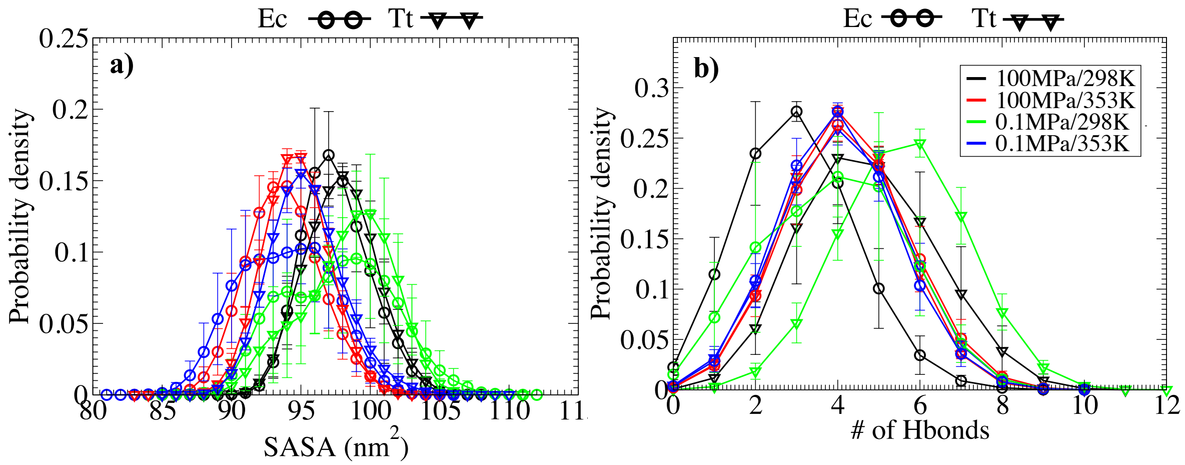

**Fig S4: a) Probability distribution for solvent accessible surface area (SASA) for both *Tt*PPase monomer (triangle) and *Ec*PPase monomer (circle); b) Number of hydrogen bonds between the residues in the catalytic pocket for both *Tt*PPase monomer (triangle) and *Ec*PPase monomer (circle). Error bars show the standard error of the mean (SEM), and those not immediately visible are at most the size of the symbol.**

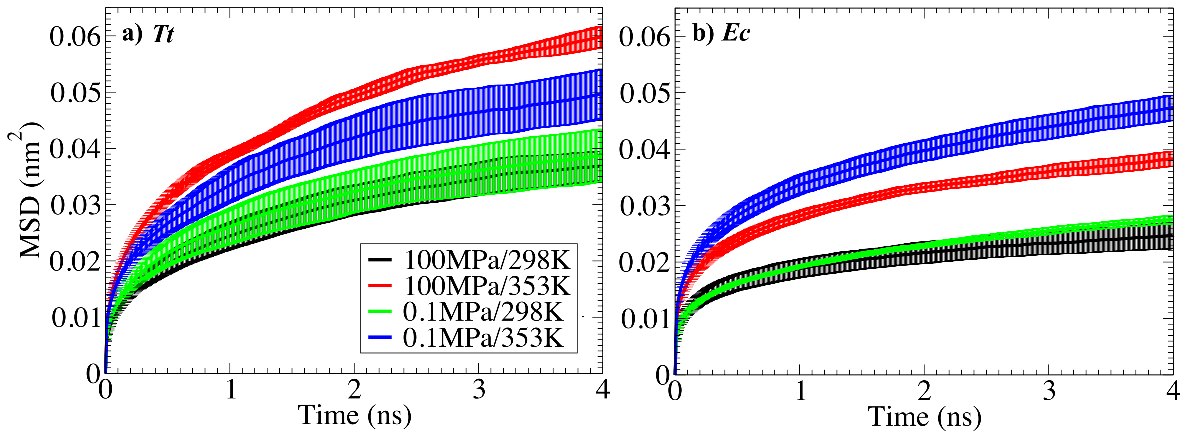


**Fig S5: Conformational flexibility determined from mean square displacement (MSD) of C-alpha atoms for (a) *Tt*PPase monomer and (b) *Ec*PPase monomer. Error bars show the standard error of the mean (SEM).**

**SI Table 1: Binding pocket residues used for the calculations**

| **Protein** | **Binding pocket residues** |
|---|---|
| ***Ec*PPase** | Asp 42,65,67,70,97,102<br>Glu 20,31<br>Arg 43<br>Lys 29, 104,142<br>Tyr 55,141 |
| ***Tt*PPase** | Asp 43,66,68,71,98,103<br>Glu 22, 32<br>Arg 44<br>Lys 30,105,141<br>Tyr 56, 140 |

**SI Table 2: Molecular dynamics simulations details**

| System type | 298K (0.1 and 100 MPa),<br>353K (0.1 and 100MPa) | Replicates |
|---|---|---|
| **Hexamer** | 500 ns | 3 |
| **Monomer** | 1000 ns | 3 |

**SI Table 3: Hexameric crystal structure comparison**

| Properties | Ec (1I6T) | Tt (3Q5V) |
|---|---|---|
| **Residues** | 1049 | 1062 |
| **Mass (KDa)** | 117.3 | 124.2 |
| **Radius of gyration (nm)** | 2.9 | 2.9 |
| **SASA ($nm^2$)** | 393 | 374 |
| **# of water in CBC** | 427 | 303 |
| **# of Hbonds between catalytic pocket residues** | 33 | 42 |
